# Effects of a single-session high-frequency repetitive magnetic stimulation on the autophagy marker LC3 and on LPS-induced inflammation in THP-1-derived macrophages

**DOI:** 10.64898/2026.04.07.716903

**Authors:** Therese B. Deramaudt, Ahmad Chehaitly, Marcel Bonay

## Abstract

High-frequency repetitive magnetic stimulation (rMS) has emerged as a non-invasive technique capable of modulating cellular signaling pathways, including those involved in inflammation and oxidative stress. Our previous work demonstrated that high-frequency rMS modulated p62/SQSTM1 expression. Given the intricate link between p62 and autophagy, we hypothesized that high-frequency rMS might influence autophagic processes in macrophages. This study investigated the effects of a single high-frequency rMS treatment on autophagy and inflammation in THP-1-derived macrophages. The results showed that 10 Hz rMS decreased autophagy, evidenced by a reduction in LC3-II expression, quantified by Western blot, and a decrease in autophagic flux, assessed by flow cytometry following bafilomycin A1 treatment. Immunofluorescence assays were used to evaluate the number of LC3-positive and LysoTracker-positive puncta. Furthermore, rMS treatment attenuated lipopolysaccharide-induced inflammation and M1 polarization in THP-1-derived macrophages, as demonstrated by the downregulation of genes encoding pro-inflammatory cytokines (IL-1β, IL-6, TNF-α) and M1 polarization markers (IL-23 and CCR7). These findings suggest that high-frequency rMS exerts a regulatory effect on autophagy and inflammation in macrophages, providing a novel approach for the treatment of inflammatory and autophagy-related diseases.

## Introduction

Repetitive magnetic stimulation (rMS) is a non-invasive technique that has gained increasing attention for its ability to modulate neuronal excitability and influence both neurological and immune functions (1–4). Initially used to treat neuropsychiatric disorders (5,6), rMS has been shown to modulate cellular signaling pathways beyond the nervous system, including those involved in inflammation and oxidative stress (7–10). The increased use of experimental animal research under controlled conditions has further enhanced our understanding and facilitated the development of clinical translational studies (11).

Recent studies have highlighted the potential of rMS to influence immune cell function, particularly in key mediators of the innate immune response, such as microglia and macrophages (9,12,13). Macrophages are highly plastic cells capable of adapting to various environmental cues and adopting distinct activation states, including pro-inflammatory M1 polarization and anti-inflammatory M2 polarization (14,15). The balance between these states is critical for effective immune responses and tissue repair, and dysregulation is linked to various pathological conditions (16). In macrophages, autophagy plays a pivotal role in maintaining cellular health and modulating immune responses by regulating inflammation and pathogen clearance (9,17,18). This process involves the formation of autophagosomes, which engulf intracellular components and deliver them to lysosomes for degradation (19).

LC3 (Microtubule-associated protein 1A/1B-light chain 3) is central to autophagy, transitioning from its cytosolic form (LC3-I) to a lipidated form (LC3-II) that associates with autophagosomal membranes. This transition serves as a marker for monitoring autophagic activity (20). While autophagy is crucial for degrading damaged organelles and misfolded proteins, the role of LC3 extends beyond cellular recycling, linking it to immune modulation (18,21). In macrophages, LC3 contributes to phagocytic mechanisms, M1/M2 polarization, and the overall immune response to pathogens (22,23). Dysregulated autophagy in immune cells has been associated with chronic inflammatory disorders, neurodegenerative diseases, and cancers, emphasizing the need to understand the regulatory mechanisms governing autophagy (24).

Our previous study demonstrated that high-frequency rMS modulates the Nrf2 signaling pathway by inducing the phosphorylation of p62/SQSTM1, a key protein involved in cellular stress responses and selective autophagy (9). Phosphorylated p62 binds to the Nrf2 inhibitor Keap1 (Kelch-like ECH-associated protein 1), facilitating the translocation of Nrf2 into the nucleus. Nrf2 is a key transcription factor that regulates antioxidant responses and plays a critical role in maintaining cellular redox balance (25,26). Given the dual role of p62 as a selective autophagy receptor and a regulator of the Nrf2 pathway, we hypothesized that high-frequency rMS could impact LC3 levels and autophagy in macrophages. Additionally, several studies have shown that autophagic pathways can modulate inflammatory signaling and immune responses. For instance, lipopolysaccharide (LPS), a potent inflammation inducer, promotes autophagy in macrophages, contributing to the degradation of intracellular pathogens and the regulation of inflammatory cytokine production (27).

In this study, we investigated the modulatory effects of high-frequency rMS on LC3-positive vesicles, pro-inflammatory cytokine expression (IL-1β, IL-6, and TNF-α), and macrophage polarization in THP-1-derived macrophages. We examined LC3-II expression levels and assessed autophagic flux following rMS treatment. Furthermore, we used LPS to induce inflammation and explored the impact of high-frequency rMS on LPS-induced inflammation and M1 polarization markers. Our findings demonstrate that high-frequency rMS reduces LC3-II levels and impairs autophagic flux, while attenuating LPS-induced pro-inflammatory and M1-related gene expression. These results suggest that high-frequency rMS modulates autophagy and inflammation in macrophages.

## Materials and Methods

### Antibodies and reagents

Maxima First Strand cDNA synthesis kit, Trizol, Fluoromount-GTM (EMS), RIPA Lysis and Extraction Buffer, sodium pyruvate, Pierce protease and phosphatase inhibitor cocktail, secondary antibodies donkey anti-mouse Alexa Fluor 488, LysoTracker red DND-99, Premo autophagy sensor GFP-LC3B kit, and iBind Western blot system were purchased from Fisher Scientific (Illkirch, France). Phorbol 12-myristate 13-acetate (PMA), dimethyl sulfoxide (DMSO), β-mercaptoethanol, para-formaldehyde (PFA), 4′,6-diamidino-2-phenylindole (DAPI), Bafilomycin A1, lipopolysaccharide (LPS), and mouse anti-GAPDH antibody were obtained from Merck (Saint-Quentin-Fallavier, France). iTaq SYBRgreen qPCR Supermix, DC protein assay kit, 4-20 % mini-Protean precast protein gels were purchased from BioRad (Marnes-la-Coquette, France).

RPMI (Roswell Park Memorial Institute) 1640 culture medium, phosphate-buffered saline (PBS), gentamicin, bovine serum albumin (BSA), and heat-inactivated fetal bovine serum were obtained from Eurobio Scientific (Les Ulis, France). Rabbit-anti-LC3 antibody was purchased from Abcam (Netherlands). IRDye^®^ 680CW goat anti-mouse IgG, and IRDye^®^ 800CW Goat anti-Rabbit IgG secondary antibodies were purchased from LI-COR^®^ Biosciences (Bad Homburg, Germany).

### High-frequency repetitive Magnetic Stimulation (rMS)

Repetitive magnetic stimulation was delivered using a B65 refrigerated butterfly coil connected to a MagPro R30 magnetic stimulator (Magventure, Denmark) purchased from Mag2Health (Villennes-sur-Seine, France). Briefly, cells were placed on the cool coil and were subjected to one session of 10 series of 100 biphasic pulses with a 25 s interval between series, resulting in a total of 1000 pulses (10 Hz at 80 % of maximum output) (9). In parallel, control cells were exposed to the same environment for the same duration as the stimulated cells but without stimulation.

### Cell culture and cell differentiation

Human monocytic THP-1 cell line (ATCC® TIB-202™) was maintained in RPMI 1640 Glutabio medium supplemented with 10 % heat-inactivated fetal bovine serum, 10 mM HEPES buffer, 1 mM sodium pyruvate, and 50 µM β-mercaptoethanol in a humidified atmosphere at 37 °C and 5 % CO_2_. Terminal differentiation of THP-1 to macrophages was obtained by washing cells twice with PBS prior to incubation with 50 nM PMA in β-mercaptoethanol-free complete RPMI 1640 medium for 48 h. After differentiation, the PMA was removed by rinsing cells with PBS and fresh complete medium was added.

### Immunoblot analysis

For extraction of total protein lysates, 1 × 10^6^ THP-1-derived macrophages were rinsed once with PBS and lyzed in ice-cold RIPA buffer supplemented with a protease and phosphatase inhibitor cocktail. After 30 min incubation on ice, cell lysates were centrifuged at 12,000 x g for 5 min at 4 °C. The supernatants containing protein extracts were collected.

Protein concentrations for total protein extracts and subcellular fractions were measured using DC protein assay kit. Protein extracts were resolved by sodium dodecyl sulfate-PAGE and transferred to a polyvinylidene difluoride membrane (Immobilon-FL, Merck). Immunoblotting was performed using the iBind flex western system according to the manufacturer’s instructions. Briefly, primary antibodies targeting LC3, GAPDH, and secondary antibodies, IRDye680RD and IRDye800RD, were diluted in iBind Flex FD solution. Fluorescence signals were acquired using Odyssey CLx imaging system (LI-COR) and densitometric analysis was achieved using Image Studio Lite v4.0.

To assess autophagic flux, cells were treated with 100 nM bafilomycin A1, which inhibits autophagosome-lysosome fusion (28).

### Total RNA isolation, reverse transcription, and real-time quantitative PCR analysis

Total RNA was isolated from treated THP-1-derived macrophages using Trizol reagent and chloroform extraction technique following the manufacturer’s instructions. RNA concentrations were determined using the NanoPhotometer® N120 (Implen; München, Germany). One µg of total RNA was reverse transcribed to cDNA using Maxima First strand cDNA synthesis kit. Quantitative analysis was achieved using real-time quantitative PCR (RT-qPCR), with each cDNA sample done in triplicate. qPCR was realized using the BioRad CFX384 Touch Real-Time PCR Detection system (BioRad) and iTaq SYBRgreen qPCR mix. Table 1 lists the specific primers used for qPCR. IL-6, IL-1β, TNF-α, IL-23, CCR7, and 18S rRNA, and were synthetized by Eurogentec (Seraing, Belgium). The cycle threshold (Ct) values of each target genes were first normalized to that of the reference gene 18S rRNA (ΔCt) then the final values (ΔΔCt values) were expressed as folds of control. Data were analyzed on the BioRad CFX manager v3.1 using the ΔΔCt method (29).

**Table 1:**
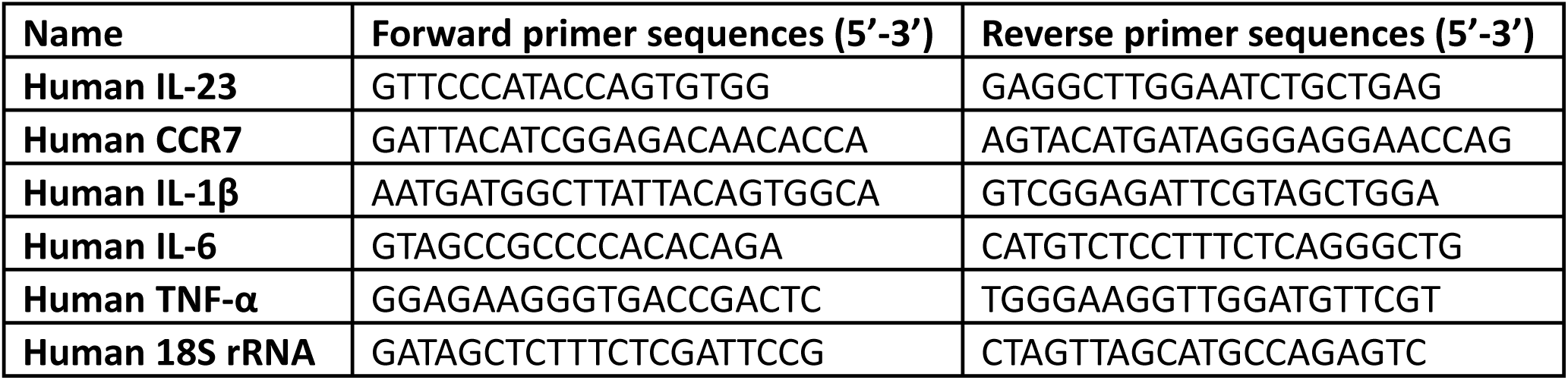
Primers used for THP-1-derived macrophages.

### Immunofluorescence microscopy

THP-1-derived macrophages were obtained by seeding 1 x 10^5^ THP-1 cells on glass coverslips in 24-well plates and differentiated with PMA for 48 h. After rMS treatment and/or LPS stimulation, macrophages were fixed in 4 % PFA in PBS, permeabilized with 0.1 % Triton X-100 for 5 min, blocked with 1 % BSA in PBS for 1 h, and incubated overnight at 4 °C with anti-LC3 antibody diluted in 1 % BSA in PBS. After washes with PBS, cells were incubated with Alexa Fluor 488-tagged secondary antibody diluted in 1 % BSA for 1 h at room temperature. For lysosomes staining, LysoTracker red DND-99 was added to live cells 1 h before the end of treatments. Nuclei were counterstained with DAPI. Coverslips were mounted on slides using Fluoromount-G. Confocal images of LC3-positive and LysoTracker red DND-99 punctate structures were acquired using Leica SP8 confocal microscope, x40 magnification. Quantitative analysis of fluorescent signals was done on 10 fields per treatment using Image J v1.53k (National Institutes of Health, Bethesda, MD, USA) and at least 100 cells per treatment were analyzed.

### Autophagy assessment and flow cytometry analysis

THP-1 monocytes were seeded at 5 x 10^5^ cells in a 6-well plates and differentiated to macrophages for 48 h with PMA. After medium change, THP-1-derived macrophages were transduced with the BacMam baculovirus included in the Premo autophagy sensor GFP-LC3B kit following the manufacturer’s recommendations. LPS 100 ng/ml and/or bafilomycin A1 treatments were added to the medium 1 h later and incubated for a total of 24 h.

Flow cytometry analysis was performed using ImageStream Mark II (Amnis, Merck) imaging flow cytometer, which allows simultaneous imaging and analysis of cells. Depending on the assay, data from 15,000 events were acquired at x40 magnification. Analyses were performed using the IDEAS v5.0 data analysis software (Amnis). Single cells were gated using a Brightfield Area versus Brightfield Aspect Ratio scatterplot, followed by Gradient RMS and Aspect Ratio versus Area to exclude debris and doublets. The GFP-LC3-positive signals were determined using the Raw Max Pixel feature on single cells at 488 nm, and gating was validated by reviewing the corresponding cell images.

### Statistical analysis

Experiments were independently performed at least 3 times. Values are presented as means ± standard errors of mean (SEM). Statistical comparisons were performed using a two-tailed unpaired t-test for comparisons between 2 groups, or one-way analysis of variance with Tukey’s multiple group comparison test for multiple comparisons. Data analyses were performed using GraphPad Prism statistical software v8.0.2. Differences were considered to be statistically significant at p < 0.05.

## Results

### High-frequency rMS decreases LC3-II expression levels in THP-1-derived macrophages

Our previous studies have demonstrated that high-frequency rMS modulates the Nrf2/Keap1 signaling pathway through increase of p62/SQSTM1 in THP-1-derived macrophages (9). Since p62 is also a well-known selective autophagy receptor involved in autophagic degradation, we sought to determine whether high-frequency rMS also modulated autophagy (30). THP-1 cells were differentiated into macrophages using PMA for 48 h before any experimental conditions were applied. To quantify LC3 expression levels, protein lysates were extracted 3 h after rMS treatment and examined by Western blot. A significant decrease in LC3-II was observed in rMS-treated THP-1-derived macrophages compared to control cells (Figure 1A). Additionally, THP-1-derived macrophages grown on coverslips were fixed 3 h after rMS treatment, and autophagosomes and lysosomes were stained with LC3 specific antibody and LysoTracker red DND-99, respectively. Confocal image analysis showed a reduction in the number of LC3-positive puncta (green puncta) in rMS-treated THP-1-derived macrophages compared to control cells (Figure 1B). Furthermore, LysoTracker red DND-99-positive puncta (red puncta) revealed a significant decrease in both the number and area of lysosomes (Figure 1B-C).

**Figure 1.**
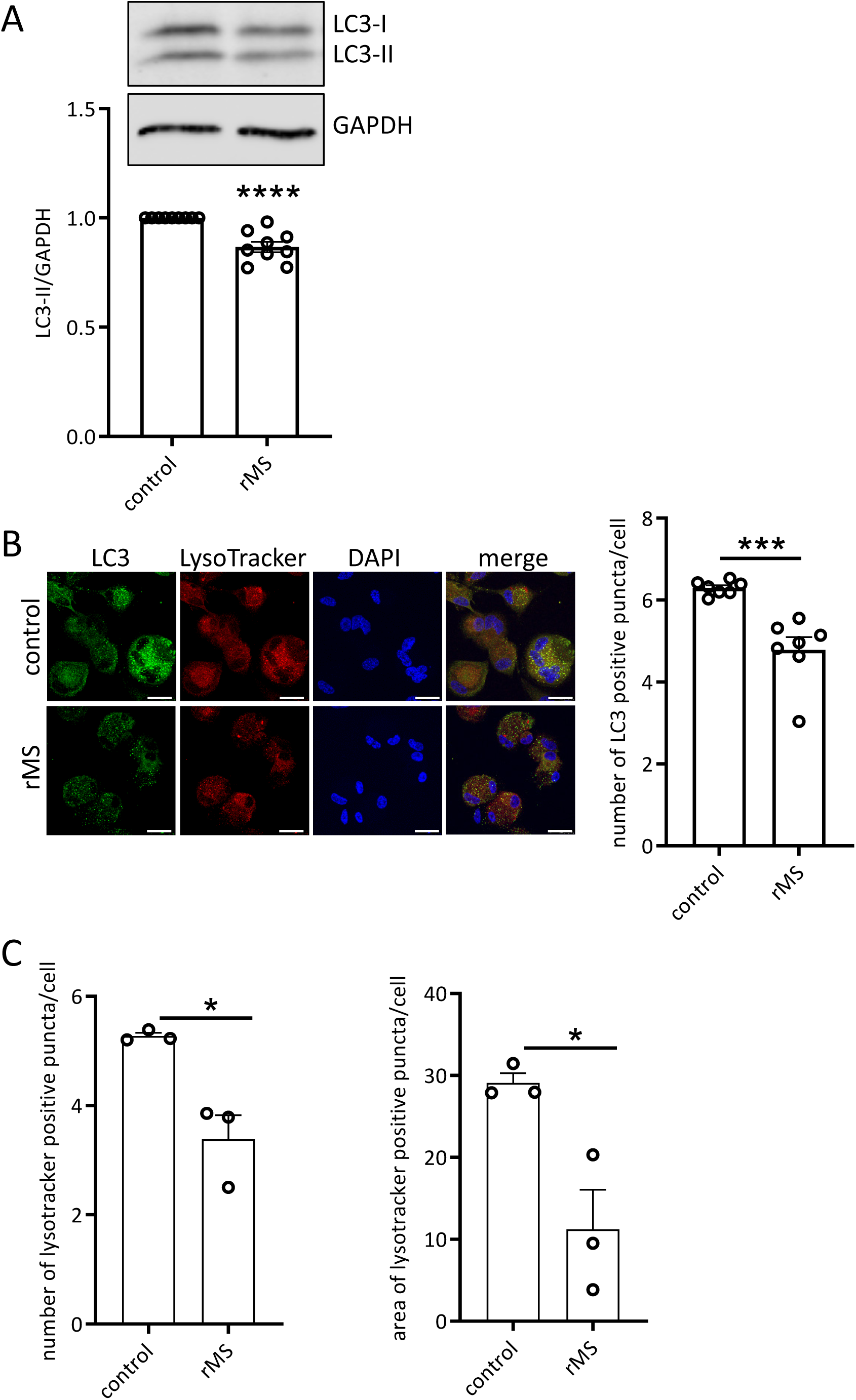
rMS reduces autophagy-lysosome pathway. (A) The autophagy-related protein LC3-I and LC3-II were assessed by Western blot and quantified using densitometry analysis. The graph on the left shows LC3-II protein levels normalized to GAPDH, while the graph on the right displays LC3-II levels normalized to LC3-I. Statistical analysis was performed using two-tailed unpaired t-test. (B) THP-1-derived macrophages treated with rMS were PFA-fixed and analyzed by confocal microscopy. Immunofluorescence images were acquired by imaging THP-1-derived macrophages. Cells were fixed 3 h after rMS stimulation and lysosomes are labeled by incubation of treated cells with LysoTracker red DND-99 1 h before cell fixation. LC3 positive puncta were obtained by labeling with antibody against LC3. Nuclei were conterstained with DAPI. White arrows indicate LysoTracker red- or LC3-positive puncta. Bars: 10 µm. Graph represents the mean number of LC3 positive puncta per cell. Analyses were based on 7 independent experiments. (C) The mean number and mean size of LysoTracker positive puncta per cell were determined using Image J. Data are shown as mean ± SEM * p < 0.05, ***p < 0.0005, ****p < 0.0001.

These data suggest a reduction in the formation of autophagosomes and lysosomes following rMS treatment.

### High-frequency rMS reduces autophagic flux

To investigate the impact of high-frequency rMS on autophagic flux, THP-1-derived macrophages were treated with 100 nM bafilomycin A1, a reversible Inhibitor of vacuolar H^+^-ATPase, which blocks autophagosome-lysosome fusion (28). After 24 h, cells were subjected to rMS treatment. Protein lysates were collected 3 h after rMS treatment and analyzed by Western blot (Figure 2A). Bafilomycin A1 treatment significantly increased LC3-II expression levels compared to control cells. In THP-1-derived macrophages treated with both bafilomycin A1 and high-frequency rMS, LC3-II protein levels also increased compared to cells treated only with high-frequency rMS. Interestingly, the rMS-induced decrease in LC3-II protein levels in bafilomycin A1-treated cells compared to control cells was evident in the LC3-II/GAPDH analysis, but not in the LC3-II/LC3-I normalization. LC3-II/GAPDH data were used to assess autophagic flux by subtracting LC3-II protein levels in untreated cells from those in bafilomycin A1-treated cells. Our data showed a significant decrease in autophagic flux in rMS-treated macrophages compared to the untreated cells (Figure 2B). Furthermore, immunofluorescence assays confirmed these results, showing a significant increase in the number of LC3-positive puncta in bafilomycin A1-treated cells compared to control cells (Figure 2C). Macrophages treated with both bafilomycin A1 and high-frequency rMS exhibited a significant decrease in the number of LC3-positive puncta compared to cells treated with bafilomycin A1 alone.

**Figure 2.**
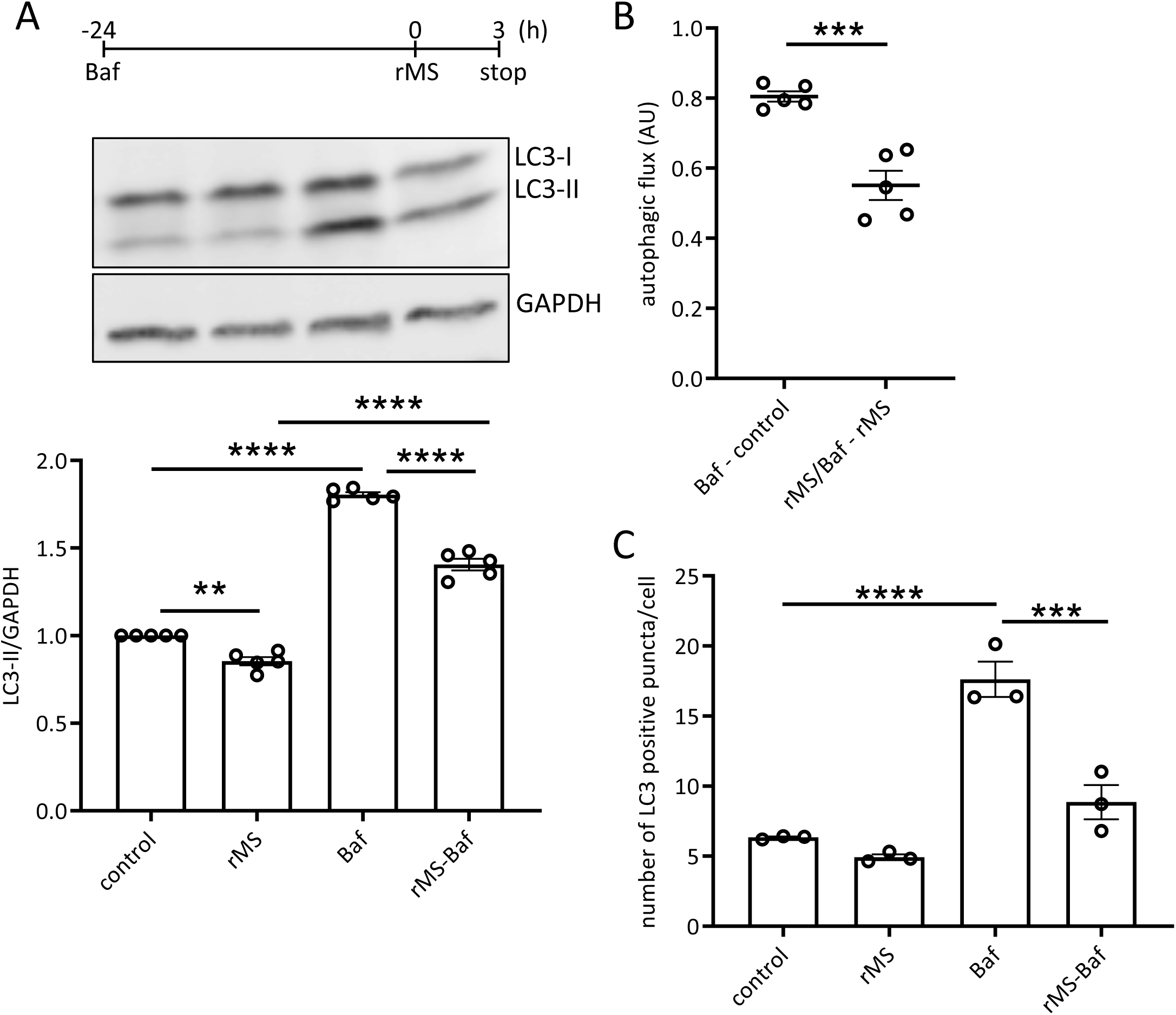
rMS reduces autophagic flux. (A) Immunoblot analysis of LC3-I and LC3-II in lysates from THP-1-derived macrophages treated with bafilomycin A1 (Baf) and stimulated with rMS. Immunoblot shown is representative of independent experiments. Data are expressed as a ratio of LC3-II and GAPDH values. (B) Autophagic flux is obtained by subtracting the LC3-II protein levels in the corresponding untreated cells from those in bafilomycin A1-treated cells (35). Data are expressed as arbitrary units (AU). (C) The number of LC3 positive puncta per cell is assessed using Image J. Data are shown as mean ± SEM ** p<0.005, *** p<0.0005, ****p < 0.0001.

Taken together, these results indicate that high-frequency rMS significantly disrupts autophagic flux.

### High-frequency rMS decreases LPS-induced inflammation

We have previously demonstrated that 10 Hz rMS treatment alone did not modulate inflammation, as indicated by the lack of induction of genes encoding the inflammatory markers IL-1β, IL-6, and TNF-α in THP-1-derived macrophages analyzed 6 h after rMS treatment (9).

To confirm the anti-inflammatory effect of high-frequency rMS in macrophages, THP-1-derived macrophages were stimulated with LPS 100 ng/ml 1 h before rMS treatment. Total RNA was extracted 6 h after LPS stimulation. RT-qPCR analysis confirmed a significant increase in mRNA expression levels of genes encoding IL-1β, IL-6, and TNF-α in response to LPS stimulation (Figure 3A). However, cells treated with rMS showed a significant reduction in the expression of LPS-induced genes, IL-1β, IL-6, and TNF-α, compared to LPS-stimulated cells alone (Figure 3B).

**Figure 3.**
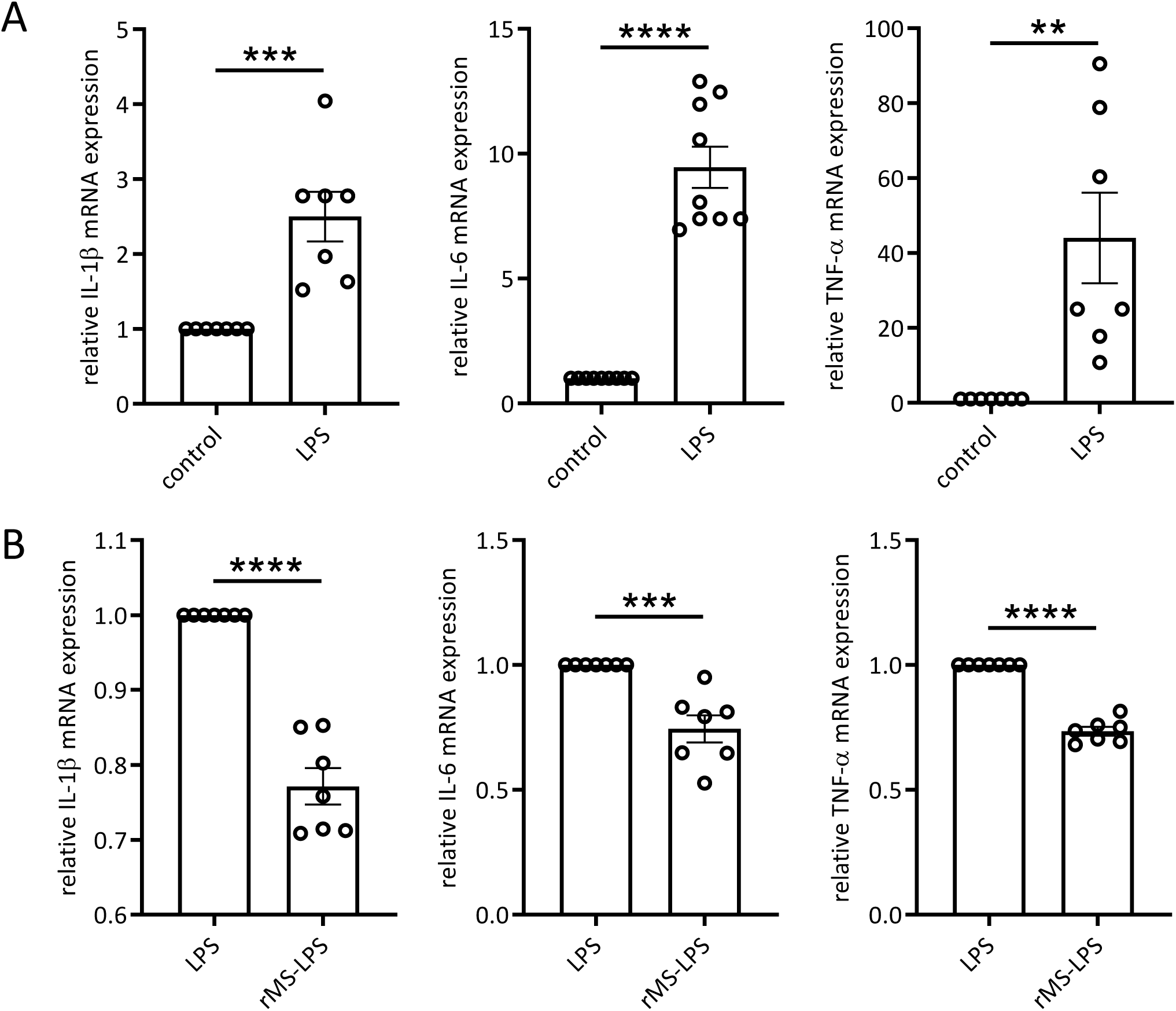
rMS decreases LPS-induced inflammation. (A) THP-1-derived macrophages were first stimulated with LPS. Total RNA was extracted from cells 6 h after LPS stimulation. Graphs show relative expression of mRNA coding for IL-1β, IL-6, and TNF-α. (B) THP-1-derived macrophages were first stimulated with LPS, then 1 h later, cells were treated or not with rMS. Total RNA was extracted from cells 6 h after LPS stimulation. Data are presented as mean ± SEM. ** p<0.005, *** p<0.0005, ****p < 0.0001.

### High-frequency rMS decreases LPS-induced M1 polarization markers

Since an increase in pro-inflammatory markers may indicate polarization of macrophages toward the M1 phenotype in response to LPS, we sought to determine whether rMS stimulation modulates LPS-induced M1 polarization. We analyzed the mRNA expression levels of two genes encoding M1 markers, IL-23 and CCR7. LPS-stimulated cells showed a significant increase in the expression of IL-23 and CCR7 compared to control cells (Figure 4A). Furthermore, rMS treatment significantly reduced LPS-induced expression levels of IL-23 and CCR7 compared to LPS-treated cells alone (Figure 4B).

**Figure 4.**
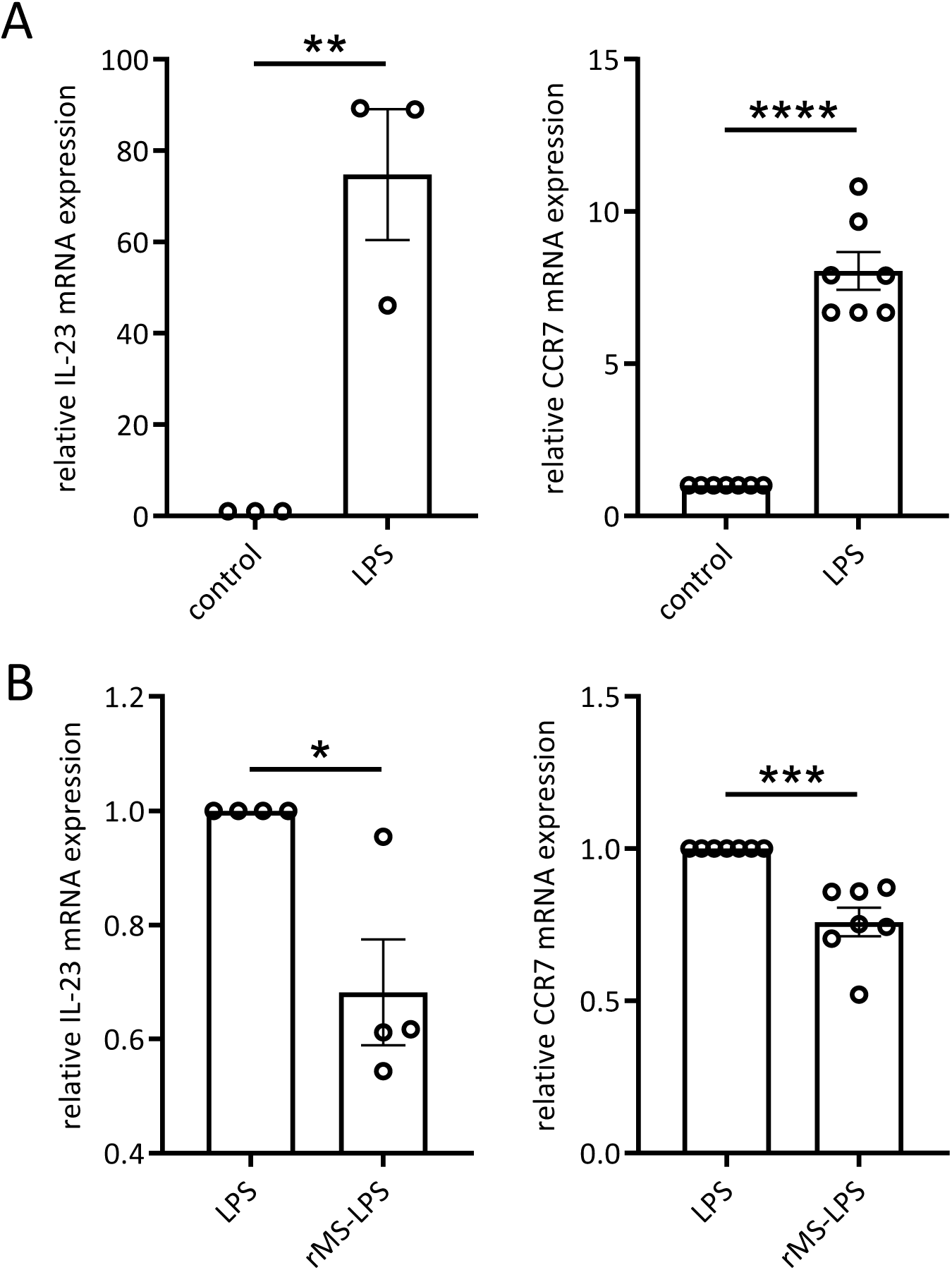
rMS treatment decreased LPS-induced M1 polarization. THP-1-derived macrophages were treated for 5 h before extraction of total RNA. Graphs show relative mRNA expression levels of genes coding for IL-23 (A) and CCR7 (B) in rMS stimulated THP-1-derived macrophages with or without presence of LPS. Data are presented as mean ± SEM. * p < 0.05, ** p<0.005, *** p<0.0005, ****p < 0.0001.

These findings suggest that rMS mitigates LPS-induced pro-inflammatory markers and M1 polarization in THP-1-derived macrophages.

### High-frequency rMS modulates autophagy independently of LPS stimulation

Previous studies have used LPS to induce autophagy in macrophages (27). To determine whether LPS stimulation could modulate autophagy in our experimental conditions, THP-1-derived macrophages were transduced with BacMam baculovirus expressing GFP-LC3, followed by the addition of bafilomycin A1 2 h later. After 24 h incubation, LPS was added 1 h prior to rMS treatment (Figure 5A). Flow cytometry was then used to analyze GFP-LC3-positive signals in THP-1-derived macrophages treated with rMS and/or LPS stimulation (Figure 5B). Analyses based on 15 000 events, from two experiments performed independently, confirmed a decrease in GFP-LC3 in rMS-treated cells compared to control cells (Figure 5C). Interestingly, LPS stimulation did not alter GFP-LC3 signals in LPS-stimulated cells compared to unstimulated cells, with or without rMS treatment.

**Figure 5.**
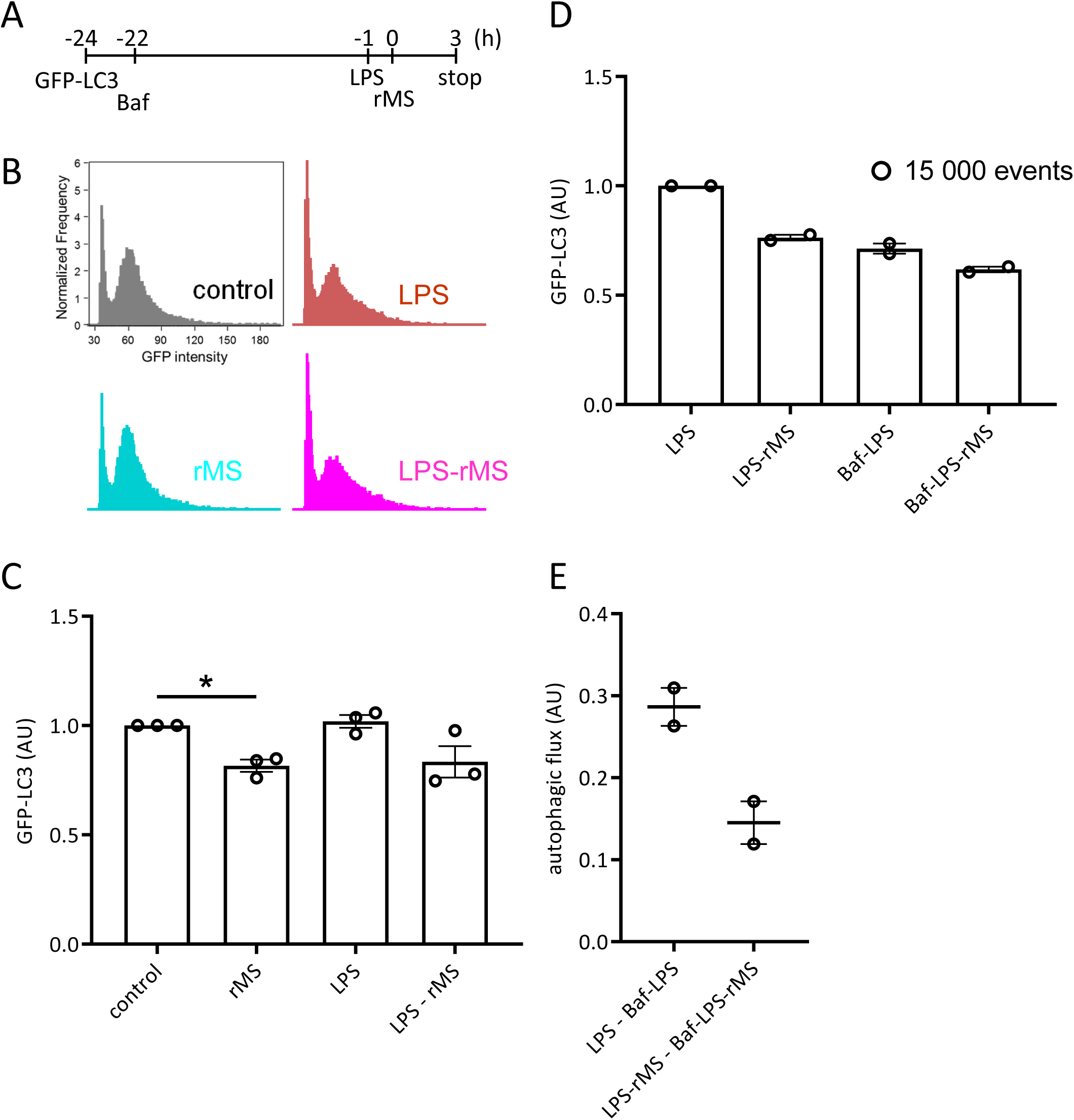
(A) Cytometry analysis of THP-1-derived macrophages are transduced with baculovirus expressing GFP-LC3. After 24 h incubation, LPS was added 1 h prior to rMS treatment. (B) Fresh cells were immediately analyzed on the flow cytometer ImageStream MRKII using IDEAS software. The GFP-LC3-positive signal is determined using the Raw Max Pixel feature on single cells at 488 nm. Representative graphs show the cell count and GFP-LC3-positive signal, and analyses were based on 15 000 events. (C) Graph represents the mean ± SEM of 3 independent experiments and data are expressed as arbitrary units (AU). * p < 0.05. (D) Bafilomycin A1 was added 2 h after transduction with the baculovirus expressing GFP-LC3. After 24 h, LPS was added 1h prior to rMS treatment. Graph shows mean ± SEM from 2 independent experiments of 15 000 events, expressed as arbitrary units (AU). (E) Autophagic flux is obtained by subtracting bafilomycin A1-treated cells to the corresponding bafilomycin A1-untreated cells.

LPS-stimulated THP1-derived macrophages pretreated with bafilomycin A1 for 24 h showed a noticeable decrease in GFP-LC3-positive signals compared to those treated with LPS alone (Figure 5D). Furthermore, the reduction in GFP-LC3 signals observed between LPS-rMS and LPS-treated cells was also evident in Baf-LPS versus Baf-LPS-rMS cells. Autophagic flux was calculated by subtracting the GFP-LC3 signal in LPS-rMS cells pretreated with bafilomycin A1 from that in LPS-rMS cells without bafilomycin A1 pretreatment (Figure 5E). A similar analysis was performed for LPS-treated cells, where the GFP-LC3 signal in cells pretreated with bafilomycin A1 was subtracted from that in untreated LPS-treated cells. The results suggest a decrease in autophagic flux in rMS-treated cells compared to those without rMS treatment.

These findings indicate that, under our experimental conditions, LPS stimulation does not impact autophagic flux in THP-1-derived macrophages. Furthermore, they suggest that rMS reduces the autophagic flux independently of LPS stimulation and also modulates LPS-induced inflammation.

## Discussion

Our findings demonstrate that high-frequency rMS is able to modulate both autophagy and inflammation in THP-1-derived macrophages, providing insight into its cellular effects relevant to immune regulation. rMS decreased autophagy activity, as indicated by reduced LC3-II protein levels, fewer autophagosomes, and decreased LC3 puncta and lysosomal content. Using bafilomycin A1 as an inhibitor of autophagosome-lysosome fusion, we confirmed that rMS impaired autophagic flux.

Our study specifically focused on LC3-II protein expression and autophagic flux, revealing that rMS treatment suppressed autophagy activity. This was evidenced by a reduction in LC3-II protein levels, fewer autophagosomes, and a decrease in LC3 puncta, as observed through confocal microscopy. Autophagic flux was assessed using LC3-II/GAPDH normalization, rather than the LC3-II/LC3-I ratio. Notably, the LC3-II/LC3-I ratio under bafilomycin A1 treatment was not significantly affected by rMS, possibly due to the saturating effect of bafilomycin A1 on autophagosome turnover, which may obscure rMS-induced alterations in LC3 processing. Moreover, since LC3-I levels were also affected by rMS, normalization to GAPDH offers a more stable reference for interpreting LC3-II changes. Autophagic flux assays further confirmed impaired autophagosome-lysosome fusion, with bafilomycin A1 validating the inhibitory effect of high-frequency rMS on autophagic flux.

This reduction in autophagic activity aligns with previous findings linking high-frequency rMS to activation of the Nrf2 pathway via increase in p62/SQSTM1 levels. Elevated levels of phosphorylated p62 enhance its interaction with Keap1, promoting Nrf2 release and nuclear translocation. Activated Nrf2 subsequently induces the expression of numerous cytoprotective and antioxidant genes, initiating anti-inflammatory and oxidative stress-inhibitory mechanisms. This pathway likely underpins the observed anti-inflammatory effects of high-frequency rMS (9). Future studies exploring the interplay between rMS, autophagy, and the p62/Keap1/Nrf2 signaling pathway will be essential to fully elucidate the effects of rMS on autophagy, oxidative stress, and inflammation.

Autophagy plays a dual role in maintaining cellular homeostasis and regulating immune responses. While its suppression by rMS can reduce inflammation, this may also hinder the survival of intracellular pathogens like *Staphylococcus aureus*, which relies on inflammatory responses for persistence (31,32). The balance between autophagic suppression and immune function underscores the complexity of rMS as a treatment.

In addition to modulating autophagy, rMS suppressed LPS-induced pro-inflammatory responses. Significant reductions in IL-1β, IL-6, and TNF-α mRNA expression, along with decreased expression of M1 polarization markers (IL-23 and CCR7), suggest that rMS mitigates macrophage polarization toward the pro-inflammatory M1 phenotype. Importantly, flow cytometry analyses showed that LPS stimulation did not alter autophagic flux under our experimental conditions, suggesting that rMS regulates inflammation through mechanisms independent of LPS-induced autophagy, potentially involving direct interference with inflammatory signaling pathways. Our future work will aim to determine whether rMS directly regulates the transcription of genes encoding pro-inflammatory cytokines or whether its effects are mediated indirectly through the activation of anti-inflammatory signaling pathways, such as Nrf2, NF-κβ, MAPK, or PI3K/Akt (8,9,33).

Our findings diverge from previous studies, such as those by Xu et al., which reported increased autophagy in human alveolar macrophages and RAW 264.7 cells following prolonged LPS exposure. The shorter LPS exposure in our study likely modulated inflammation without significantly affecting autophagy (27). Specifically, macrophages in our study were exposed to LPS for only a few hours, in contrast to the 24 h exposure used in their study. This shorter exposure time appears sufficient to modulate inflammation with LPS but not to affect autophagy to a level detectable by flow cytometry.

Similarly, Wang et al. demonstrated enhanced autophagic flux in bone mesenchymal stromal cells treated with high-frequency rTMS (34). They used a 50 Hz stimulation protocol, consisting of 20 trains of 100 pulses daily for 3 to 5 days, resulting in increased LC3-II protein levels, enhanced autophagic flux, and decreased p62 levels after repeated rTMS treatments. They proposed the involvement of the NMDAR-Ca2^+^-ERK-mTOR signaling pathway in BMSCs. Discrepancies between these findings and ours may arise from differences in experimental conditions, including stimulation frequency, pulse intensity, and treatment duration. In our experiments, 10 Hz rMS induced significant effects on the Nrf2 pathway, whereas 5 Hz and 20 Hz showed minimal impact (data not shown). Further research is required to determine how treatment parameters, such as stimulation frequency, pulse intensity, and treatment duration, influence rMS outcomes across different cell types and experimental contexts. Additionally, signaling pathways involved in modulating inflammation and macrophage function remain to be explored. Further research is required to determine how treatment parameters, such as stimulation frequency, pulse intensity, and treatment duration, influence rMS outcomes across different cell types and experimental contexts. Additionally, alternative pathways involved in modulating inflammation and macrophage function remain to be explored. While many studies on repetitive magnetic stimulation focus on excitable cells, our model uses non-excitable THP-1-derived macrophages. In this context, biophysical factors such as energy deposition, electric field strength, and frequency-dependent modulation of membrane receptors or ion channels may alter calcium homeostasis and signaling pathways such as MAPK, PI3K/Akt, AMPK. These can ultimately influence autophagy in a cell-type and stimulus-dependent manner.

Our study highlights the dual effects of rMS on autophagy and inflammation, suggesting its potential value for inflammatory and autoimmune diseases. By suppressing both autophagic flux and pro-inflammatory markers, rMS could help restore cellular homeostasis in conditions characterized by dysregulated immune responses, such as neurological and inflammatory disorders. Additionally, combining rMS with strategies to activate Nrf2 may enhance its efficacy by synergistically reducing inflammation and oxidative stress.

However, reduced autophagic flux may lead to the accumulation of cellular debris and damaged organelles, posing a risk of long-term cellular dysfunction. A careful modulation of rMS parameters will be necessary to balance the benefits of reduced inflammation with the need for adequate cellular clearance mechanisms. Future studies should also investigate the precise molecular pathways through which rMS influences autophagy and inflammation, including its effects on key regulatory proteins such as Keap1, p62, and LC3, as well as the potential involvement of autophagy in modulating the Nrf2/Keap1/p62 signaling pathway. Expanding this research to *in vivo* models will be crucial for validating its potential. Optimizing treatment parameters, including stimulation frequency, pulse intensity, and treatment duration, will also enhance its translational applicability.

In conclusion, this study provides the first evidence that high-frequency rMS reduces autophagy and inflammation in THP-1-derived macrophages via autophagic modulation. These findings provide valuable insights into the cellular mechanisms underlying effects of rMS.

## Study limitations

Our study examined only a single session of high-frequency (10 Hz) rMS in THP-1-derived macrophages. While this protocol reduced autophagy and LPS-induced inflammation/M1 polarization, the optimal parameters for these effects remain unknown. Future studies should consider varying the frequency, pulse intensity, number of pulses, and treatment duration to define the optimal conditions for modulating specific cell types.

Discrepancies with prior reports of rMS-induced autophagy may reflect differences in stimulation protocols and cell types, particularly between non-excitable macrophages and excitable cells. Additionally, the signaling pathways linking rMS to suppressed autophagic flux and attenuated inflammation, despite the typical anti-inflammatory role of autophagy, require further elucidation.

## Acknowledgments

We acknowledge the technical support of the imaging core facility located in UFR des Sciences de la Santé, UVSQ.

## Author contributions

T.B.D.: Conceptualization, Investigation, Methodology, Supervision, Validation, Formal analysis, Visualization, Writing – original draft, Writing – review and editing. A.C.: Methodology, review. M.B: Conceptualization, Supervision, Funding acquisition, Validation, Writing – review and editing. The authors declare that all data were generated in-house.

## Data Availability Statement

The data reported in this study is available from the corresponding author upon request.

## Abbreviations

CCR7: C-C motif chemokine receptor 7
cDNA: complementary deoxyribonucleic acid
DAPI: 4′,6-diamidino-2-phenylindole
IL-1β: interleukin-1 β IL-6 interleukin-6
IL-23: interleukin-23
Keap1: Kelch-like ECH-associated protein 1
LC3: Microtubule-associated protein 1A/1B-light chain 3
LPS: lipopolysaccharide
Nrf2: nuclear factor erythroid 2–related factor 2
p62/SQSTM1: sequestosome 1
PBS: phosphate-buffered saline
PFA: paraformaldehyde
RIPA: radioimmunoprecipitation assay buffer
rMS: repetitive magnetic stimulation
RNA: ribonucleic acid
RT-qPCR: reverse transcription-quantitative polymerase chain reaction
THP-1: human leukemic cell line
TNF-α: tumor necrosis factor-α

